# Large-scale *in-vitro* production of red blood cells from human peripheral blood mononuclear cells

**DOI:** 10.1101/659862

**Authors:** Steven Heshusius, Esther Heideveld, Patrick Burger, Marijke Thiel-Valkhof, Erica Sellink, Eszter Varga, Elina Ovchynnikova, Anna Visser, Joost H.A. Martens, Marieke von Lindern, Emile van den Akker

## Abstract

Transfusion of donor-derived red blood cells is the most common form of cellular therapy. Donor availability and the potential risk of alloimmunization and other transfusion-related complications may, however, limit the availability of transfusion units especially for chronically transfused patients. *In-vitro* cultured, customizable red blood cells would negate these concerns and introduce precision medicine. Large-scale, cost effective production depends on optimization of culture conditions. We developed a defined medium and adapted our protocols to GMP culture requirements, which reproducibly provided pure erythroid cultures from peripheral blood mononuclear cells without prior CD34^+^ isolation, and a 3×10^7^-fold increase in erythroblasts in 25 days. Expanded erythroblast cultures could be differentiated to CD71^dim^CD235a^+^CD44^+^CD117^−^DRAQ5^−^ red blood cells in 12 days. More than 90% of the cells enucleated and expressed adult hemoglobin as well as the correct blood group antigens. Deformability and oxygen binding capacity of cultured red blood cells was comparable to *in-vivo* reticulocytes. Daily RNA sampling during differentiation followed by RNA-seq provided a high-resolution map/resource of changes occurring during terminal erythropoiesis. The culture process was compatible with upscaling using a G-Rex bioreactor with a capacity of 1L per reactor, allowing transition towards clinical studies and small-scale applications.

## Introduction

Blood transfusion is the most applied cellular therapy, with more than 80 million transfusion units administered worldwide each year^1^. Inherent risks of donor-transfusion material are alloimmunization and presence of blood-borne diseases. Oxygen-carrier substitutes have shown to be applicable in case of immediate emergency but cannot replace long term blood transfusions^2^. The potential to culture red blood cells (RBC) for transfusion purposes has been recognized for a long time^3–10^. Cultured RBC (cRBC) that are antigen-compatible will decrease the risk of alloimmunization in patients. Cost-effective, large-scale culture of blood group matched RBC will provide a degree of donor independency and minimization of donor-patient blood type variation. In addition, cRBC can be used as vehicles for enzyme replacement therapy^11^, or as therapeutic delivery systems targeting specific body parts^12^. Several groups already cultured enucleated cRBC from cord blood CD34^+^cells^13–15^. However, these cells produce fetal hemoglobin with a higher tendency to denature and to cause membrane damage compared to adult hemoglobin^16^. We have previously shown that enucleated cRBC can be generated starting from adult peripheral blood mononuclear cells (PBMC), a better accessible source than cord blood CD34^+^ cells and allows adult autologous cRBC^17^. Importantly, the erythroid yield from PBMC is 10-15 fold increased compared to CD34^+^ cells isolated from a similar amount of PBMC, due to support from CD14^+^ cells present in PBMC^17–19^.

One transfusion unit contains about 2×10^12^ RBC, reflecting the high requirement for erythroblast expansion to obtain sufficient numbers of cRBC. Previous attempts to culture the required number of enucleated cRBC from CD34^+^ cells isolated from PBMC were hampered by low expansion or poor enucleation^20,21^. Expansion of CD71^high^CD235a^dim^ erythroblasts can be prolonged by exploiting the cooperative action of erythropoietin (EPO), stem cell factor (SCF) and glucocorticoids involved in stress-erythropoiesis in a serum/plasma-free environment^7,17,18,22,23^, while differentiation is induced by increasing concentrations of EPO and dispensing SCF and glucocorticoids. Here, we describe a three stage GMP-grade culture protocol using culture dishes or G-Rex bioreactors, both with high expansion and enucleation to generate PBMC-derived cRBC. To this end we have developed a completely defined GMP-grade medium. This three-stage culture protocol can be used for small-scale GMP-grade production, yielding >90% enucleated reticulocytes with adult hemoglobinization.

## Material and Methods

### Cell culture

Human PBMC were purified by density separation using Ficoll-Paque (manufacturer protocol). Informed consent was given in accordance with the Declaration of Helsinki and Dutch National and Sanquin Internal Ethic Boards. PBMC were seeded at 5-10×10^6^ cells/ml (CASY^®^ Model TCC; Schärfe System GmbH, Reutlingen, Germany) in Cellquin medium based on HEMA-Def^7,17^ with significant modification (Table S1 lists all components) supplemented with erythropoietin (2U/ml; ProSpec, East Brunswick, NJ), human recombinant stem cell factor (100ng/ml; ITK Diagnostics BV, Uithoorn, The Netherlands), dexamethasone (1μM; Sigma, St. Louis, MO) and 0.1% human serum albumin (cHSA; kindly provided by Sanquin Plasma Products, Amsterdam, The Netherlands, perturbation with albumin see Supplementary Material), referred to as Expansion Medium (EM). Interleukin 3 (IL3) was added (1ng/ml; Miltenyi Biotec, Bergisch Gladbach, Germany) to EM on the first day (stage I). Media was partially replenished every two days with EM. Around day 6, upon erythroblasts detection, the cells were maintained 1-2×10^6^ cells/ml for 15-25 days (stage II). Erythroblasts differentiation (stage III) was induced in Differentiation Medium (DM) containing Cellquin supplemented with erythropoietin (10U/ml), 5% Omniplasma (Omnipharma S.A.L., Beirut, Lebanon), holotransferrin (700μg/ml; Sanquin) and heparin (5U/ml; LEO Pharma BV, Breda, The Netherlands). At day two, half a medium change was performed. At day 5, storage components (Sanquin Plasma Products, Amsterdam, The Netherlands) were added and media was refreshed (half) every two days until fully differentiated at day 10-12 of differentiation. For reticulocyte filtration see Supplementary Material.

### Flow cytometry

Cells were washed and resuspended in HEPES buffer supplemented with 1% HSA. Cells were incubated with primary antibodies for 30 minutes at 4 °C, measured on FACS Canto II or LSRFortessa (both BD Biosciences, Oxford, UK) and analyzed using FlowJo software (FlowJo v1O, Ashland, OR, USA). Reticulocyte RNA was stained with Thiazole orange (TO; Sigma) as described previously^24^ and DNA using DRAQ5 (1:2500; Abeam, Cambridge, UK). Antibodies are listed in Supplementary Material.

### RBC deformability

RBC deformability was measured by the Automated Rheoscope and Cell Analyzer (ARCA) as described previously^25^. 10 Pa shear stress was used and 3000 cells were measured and grouped in 30 bins according to increasing elongation or cell projection area (as a measure of membrane surface area). Both the extent of elongation (major cell radius divided by minor cell radius) and the area (in mm^2^) was plotted against the normalized frequency of occurrence.

### HPLC

Culture lysates were prepared and stored at −80 °C prior to analysis as described previously^25^. In short, hemoglobin separation was performed by high performance cation exchange liquid chromatography (HPLC) on Waters Alliance 2690 equipment (Waters, Milford, MA, USA) using30min elution over a combined 20-200mM NaCl and pH7.0-6.6 gradient in 20mM BisTris/HCl and 2mM KCN. A PolyCAT A 100/4.6mm, 3mm, 1500Å column (PolyLC, Columbia, MD, USA).

### Coomassie

Ghosts (RBC membranes) were generated as described before^26^ subjected to SDS-PAGE gels (Bio-Rad, Hercules, CA, USA) and total proteins stained with Coomassie. In short, proteins were fixed in 30% ethanol, 2% (v/v) phosphoric acid overnight. Washed two times 10minutes in 2% phosphoric acid and equilibrate for 30min in 2% phosphoric acid containing 18% ethanol and 15% (w/v) ammonium sulphate. Stain by diluting Coomassie Blue G-250 slowly in water from 0.2% to a final concentration of 0.02% (0.2mg/ml).

### Cytospins

Cells were cytospun using Shandon Cytospin II (Thermo Scientific), dried and fixed in methanol. Cells were stained with benzidine in combination with the Differential Quick Stain Kit (PolySciences, Warrington, PA) (manufacturers protocol). Slides were dried, subsequently embedded in Entellan (Merck-Millipore) and covered with a coverslip. Images were taken using microscope DM2500 with 40x or 10x object (Leica DM-2500; Germany).

### RNA-seq analysis

Erythroid cultures on differentiation medium were sampled daily for 12 days. Sequencing libraries were prepared using Trizol RNA isolation, cDNA amplification and rRNA depletion using HyperPrep Kit with RiboErase (KAPA Biosystems, Pleasanton, CA, USA) as described by manufactures. Reads were mapped to ChGR38.v85 and differential expression analysis was performed with EdgeR^27^. A detailed description can be found in Supplementary Material.

## Results

### Large-scale erythroblast expansion from PBMC using a G-Rex bioreactor and GMP-grade medium

To establish medium conditions to obtain and prolong erythroid expansion from adult PBMC, we first tested distinct sources of albumin. Different expansion media (EM, Table S1) with 0.1% ultra-clean human serum albumin obtained after plasma fractionation (cHSA), 2,5% plasma, 2,5% Albuman^®^, 0.1% detoxified HSA (dHSA), or 0.1% recombinant HSA (rHSA) were used (Figure 1A and Figure S1A). Plasma or Albuman^®^ resulted in (i) low erythroblast yield, (ii) presence of non-erythroid cells (negative for CD71 and CD235) and (iii) premature differentiation of erythroblasts, indicated by a loss of CD71 expression in conjunction with increased CD235a expression (Figure S1A)^28,29^. In contrast, EM supplemented with cHSA, dHSA, or rHSA showed a significantly increased erythroid expansion potential with limited spontaneous differentiation and a complete absence of non-erythroid cells (Figure 1A and Figure S1A). Of note, PBMC contain primarily T-cells, myeloid cells and B-cells and only on average 0.16% CD34^+^ HSPC that are capable of differentiating into erythroid cells^17,19^. Therefore, expansion curves using PBMC as a starting material show a drop in expansion between day 0 and day 5, caused by a loss of these non-proliferating immune effector cells^17,18^.

**Figure 1.**
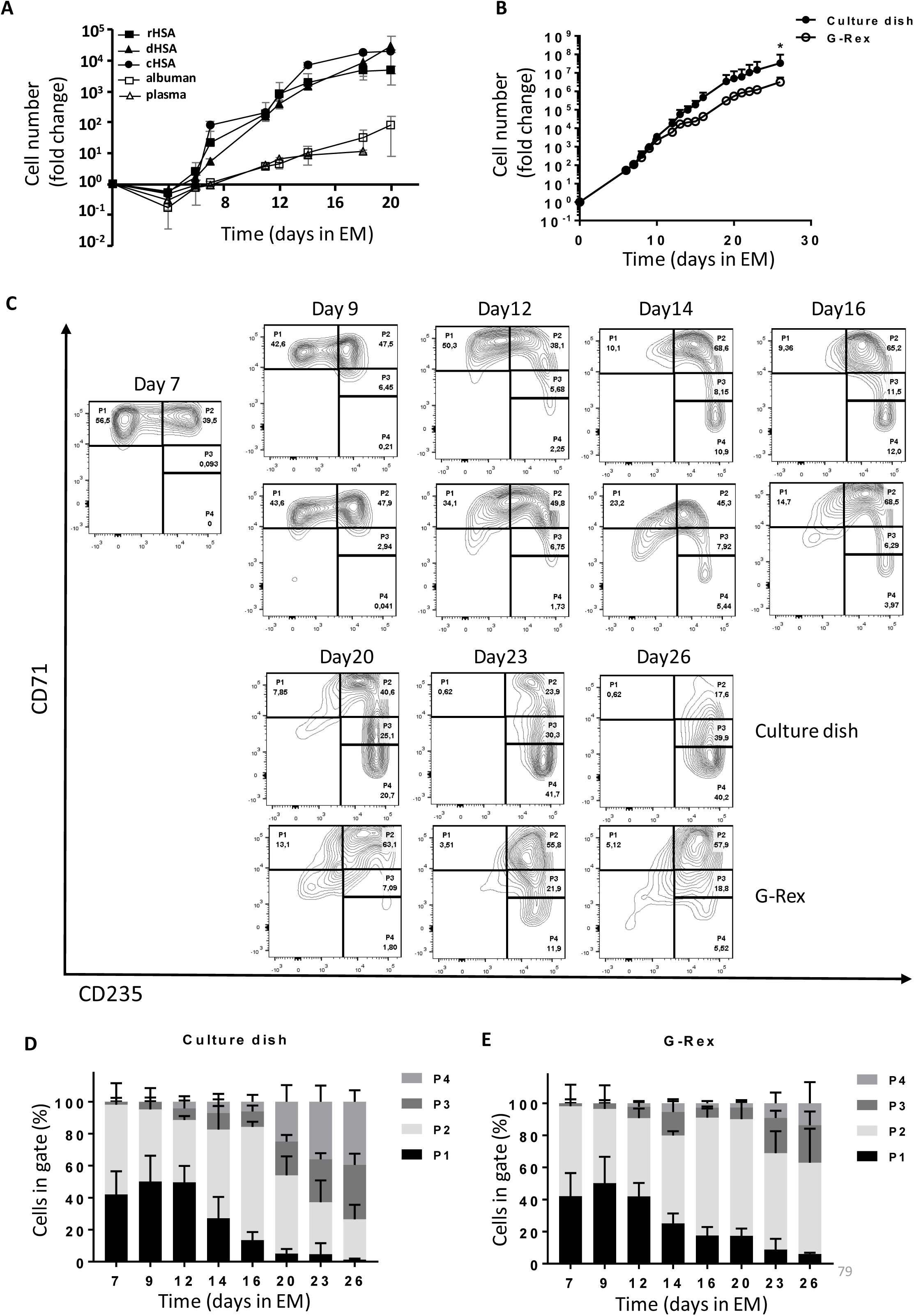
Efficient expansion of erythroblasts in plasma/serum-free GMP-grade medium. **(A)** PBMC were cultured towards erythroblasts in GMP-grade medium supplemented with EPO, SCF and Dex (EM) for 26 days. Medium was prepared using either ultra-clean HSA (cHSA), detoxified HSA (dHSA), recombinant HAS (rHSA), Albuman^®^ or plasma. Cell counts at day 0 were normalized to 1 PBMC at the start of culture. Cultures were kept at 0.7 to 2×106 by dilution. Symbols indicate mean fold-change at any day compared to 1 PBMC seeded, error bars indicate SD (n=4). **(B)** PBMC from 4 independent donors were cultured from PBMC in Cellquin medium (cHSA) supplemented with EPO (2U/ml), SCF (100ng/ml) and Dex (1μM) in culture dishes until a pure erythroblast population was obtained (at day 7). Erythroblasts were further expanded in a G-Rex bioreactor or in culture dishes. Mean fold-change (± SD) was calculated and compared (two-way ANOVA, * *P*<0.05; n=4). **(C)** Representative dot plots indicating cell surface expression levels of CD71 and CD235 in cultures as described in **B**. Quadrants are labeled (P1-P4) and relative cell numbers per quadrant indicated as percentage **(D-E)** quantification of percentages per quadrant in dot blots similar to those shown in C. Error bars indicate SD (n=4) D: cells cultured in dishes. E: cells cultured in G-Rex

Large scale cRBC production in culture dishes is practically impossible. Therefore, a G-Rex bioreactor from Wilson Wolf Manufacturing (Saint Paul, MN, USA) was used in which a gas-permeable membrane at the bottom allowed proliferation in larger volume/surface conditions^30,31^. The G-Rex bioreactor does not support adherent cells, such as the CD14^+^ PBMC that promote erythroid yield by increasing CD34^+^ cell survival^18^, which compromises stage I yield (data not shown). Therefore, PBMC cultures were started in culture dishes/cell stacks until an erythroblast population was obtained around day 7 of culture in EM supplemented with cHSA. Subsequently (stage II), cultured erythroblasts were either transferred G-Rex systems or retained in culture dishes and could be maintained for at least 26 days (Figure 1B). Transfer to a G-Rex bioreactor briefly delayed erythroid expansion but showed a similar expansion rate between day 15 and 25. Although the G-Rex bioreactor yielded slightly less cells (3×10^6^ *vs.* 3×10^7^), less donor variation was observed. Erythroid cells can be staged from CD71^high^CD235^dim^ erythroblasts (P1) to enucleated CD71^−^CD235^+^ reticulocytes (P4), with intermediate stages in which cells are characterized by increased CD235a and reduced CD71 expression (Figure 1C)^29^. At day 7, no non-erythroid cells (CD71^−^CD235^−^) were observed, indicating a pure erythroid culture, (day 0 to day 7 culture progression was previously published by our laboratory^18^). During erythroblast expansion in stage II, erythroblasts gained expression of CD235a from day 10 onwards resulting in a pure population of CD71^high^CD235^+^ erythroblasts, that could be expanded for at least 26 days (Figure 1C-E and Figure S1B). Interestingly, cells expanded in the G-Rex system maintained a less differentiated state for a prolonged culture period (>16days), as shown by delayed CD235a upregulation and maintenance of high CD71 expression (Figure 1E). In conclusion, a defined GMP-grade medium enabled pure erythroblast cultures from a mixed PBMC cell pool with significantly delayed onset of spontaneous differentiation, resulting in a large expansion potential in both culture dishes and a G-Rex bioreactor.

### Erythroid cultures from G-Rex and normal culture dishes fully enucleate

Erythroblast differentiation (stage III) is induced by removing SCF and dexamethasone whilst increasing the EPO and holo-transferrin concentration and supplementing with 5% Omniplasma. Heparin is added to prevent medium from clotting (differentiation medium, DM). Erythroblast differentiation is characterized by i) a transient proliferation burst with decreased cell-cycle time resulting in a reduced cell volume, ii) hemoglobinization, iii) erythroid specific protein expression of e.g., blood group antigens, and iv) enucleation^13,17,28,29,32^. During the first days of differentiation, in particular in culture dishes, proliferation was observed followed by cell growth arrest (Figure 2A). Note that the number of cells after the initial short proliferation burst did not decrease, suggesting no negative effect on culture viability. In contrast, cells differentiated in a G-Rex bioreactor did not show increased expansion. During erythroblast differentiation, both erythroid cells cultured in dishes and the G-Rex system increased expression of CD235a and lost expression of CD71, although erythroblasts differentiated slightly faster in culture dishes (Figure 2B-D). Differentiation was accompanied with a decrease in cell size (FSC-A). Both CD71 and c-KIT (CD117) expression increased at day 1 followed by a sharp down regulation (Figure S2A-B). Furthermore, CD44 was progressively reduced in expression as reported before^17^. Although CD235 expression increases in early differentiation as cells become smaller it decreases slightly, which may be due to loss of membrane surface during enucleation.

**Figure 2.**
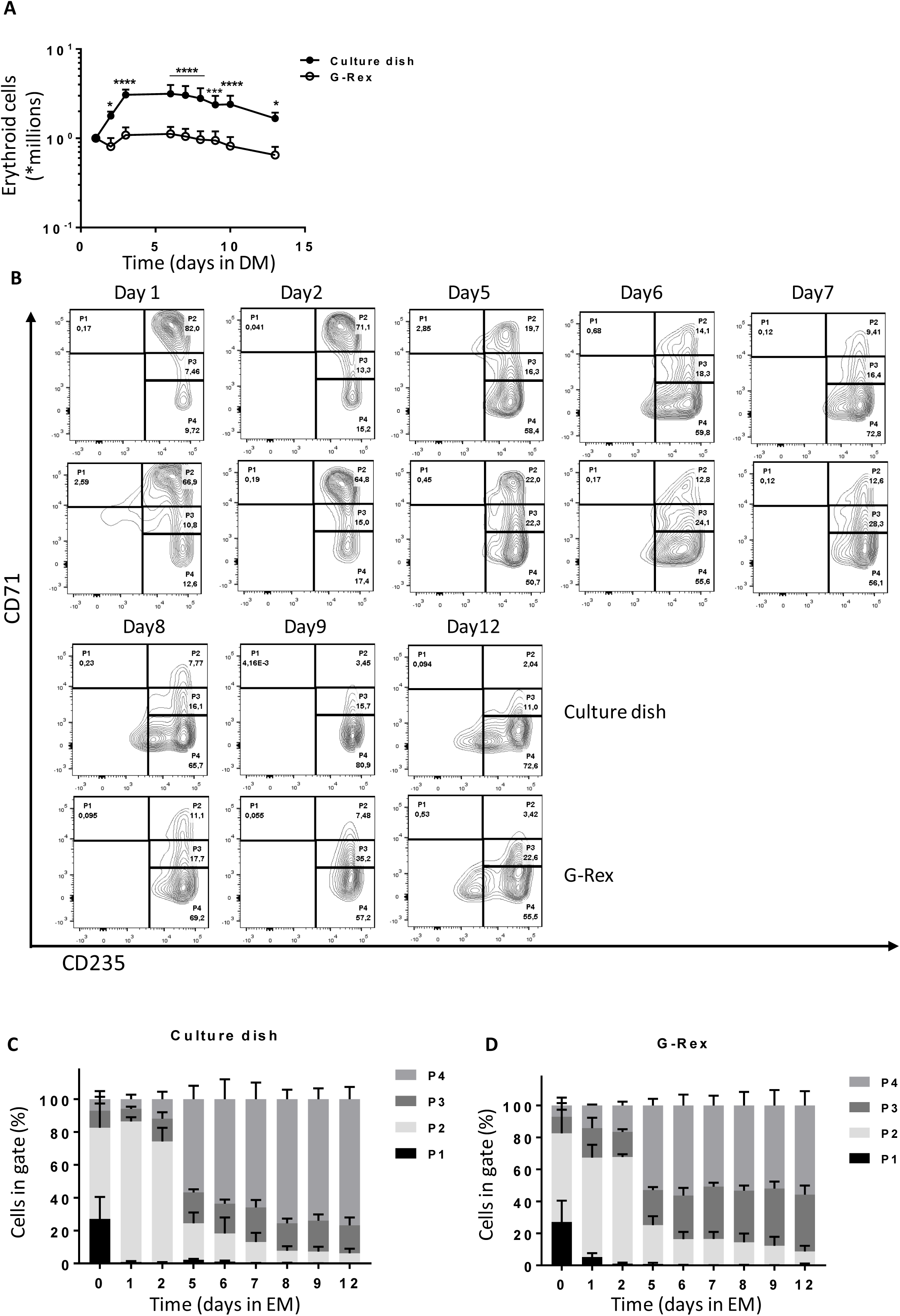
Differentiation of erythroblasts in culture dishes or a G-Rex bioreactor. **(A)** Erythroblast cultures were established in culture dishes. Erythroblast were washed and reseeded at 1×10^6^/ml in Cellquin medium supplemented with EPO (10U/ml), Transferrin (700μg/ml), 5% human plasma, and heparin (5U/ml) (Differentiation medium, DM) in culture dishes (closed symbol) or G-Rex (open symbol). Erythroblasts were differentiated for 12 days. Cell density was measured at days indicated. Mean cell numbers were calculated. Error bars indicate SD Cell expansion was compared by two-way ANOVA, \**P*<0.05, \*\*\**P*<0.001, \*\*\*\**P*<0.0001; n=4. **(B)** Representative dot plots indicating cell surface expression levels of CD71 and CD235 in cultures as described in **A.** Quadrants are labeled (P1-P4) and relative cell numbers per quadrant indicated as percentage **(C-D)** quantification of percentages per quadrant in dot blots similar to those shown in C. Error bars indicate SD (n=4) C: cells cultured in dishes. D: cells cultured in G-Rex

### Factors that affect enucleation at the end of erythroid differentiation

At blood banks, RBC are stored in media that contain specific components protecting viability, which were not present in DM. Therefore, a specific storage component solution was added at day 5 of differentiation, when the first reticulocytes arose in culture. This increased the number and frequency of enucleated cRBC, particularly in the G-Rex bioreactor (data not shown). Nonetheless, initial differentiation experiments in the G-Rex system yielded low numbers of enucleated cells compared to culture dishes (Figure S3A). Importantly, EM is completely replaced by DM upon initiating of differentiation in dishes, whereas only 90% EM could be replaced with DM in the G-Rex system. Indeed, replacing 90% of the culture medium in dishes, resulted in a reduction of enucleation, compared to complete medium replacement (56% vs 85%; Figure S3B). Reduction of dexamethasone (from 1μM to 10nM) two days prior to differentiation induction did not affect enucleation (Figure S3D). However, SCF removal from the culture medium two days prior to differentiation in culture dishes increased enucleation 1.5-fold (56% *vs.* 85%; Figure S3C). This indicated that residual SCF at the start of differentiation negatively affects enucleation during terminal differentiation. Enucleation was observed from day 5 onwards, resulting in >90% enucleation after 12 days of differentiation in culture dishes and almost 85% enucleation in the G-Rex system (Figure 3A and Figure S3E). A slight difference in the ratio nuclei/cRBC between culture dishes and a G-Rex bioreactor was observed (Figure 3B). The flow cytometry data and cytospin images at the end of differentiation revealed pyrenocytes (nuclei encapsulated by plasma membrane) and some nucleated cells (Figure 3C and Figure S3E). Filtration using neonatal leucoreduction filters resulted in 99% removal of nuclei and remaining nucleated cells and yielded a homogenous population of enucleated cRBC (Figure 3C).

**Figure 3.**
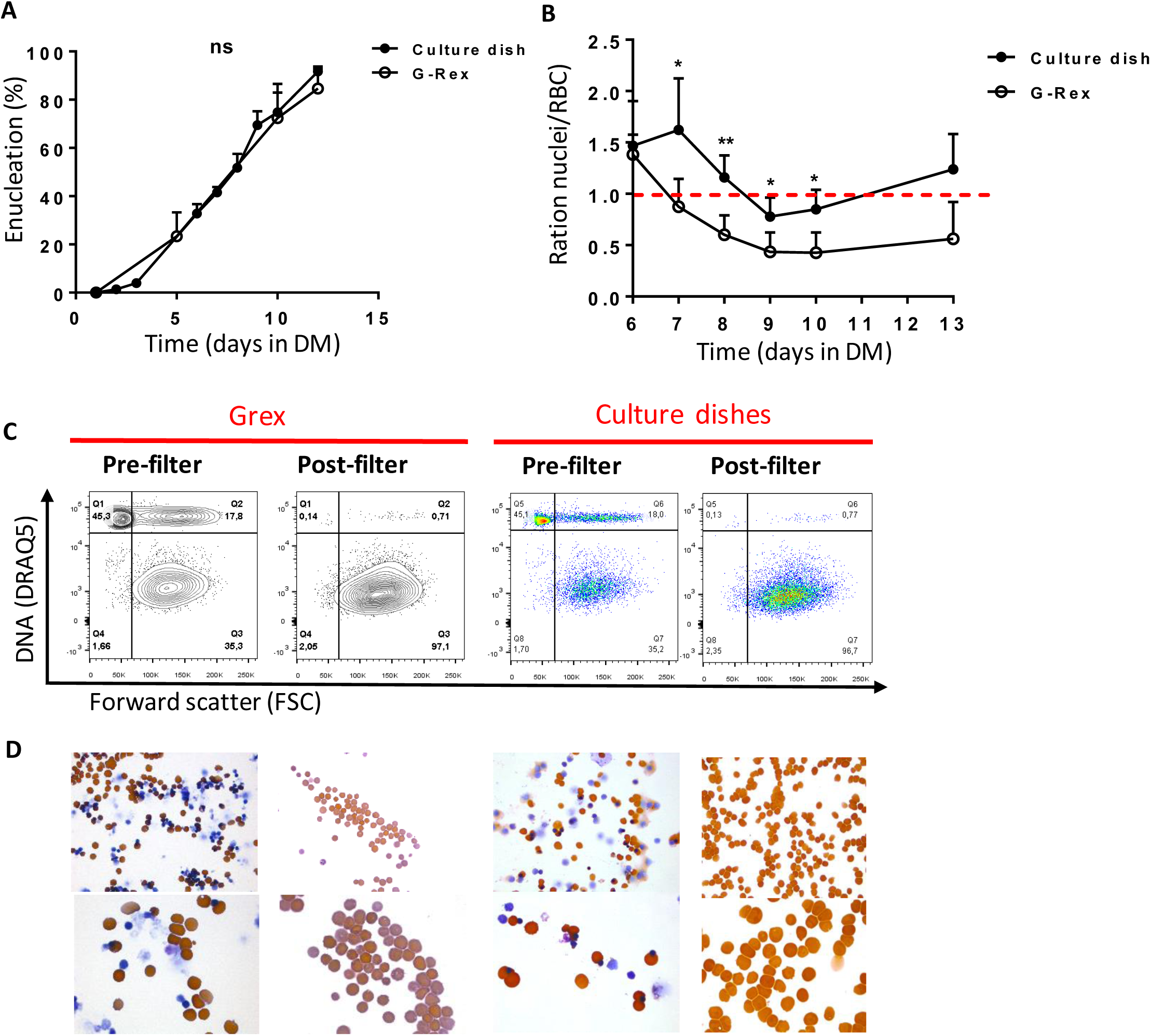
Efficient enucleation is observed after 10 days in differentiation medium. **(A-C)** enucleated cells and nuclei were measured by flow cytometry, using DRAQ5 DNA staining against size (forward scatter) as shown in **C. (A)** The percentage of enucleated erythroid cells was measured during differentiation in culture dishes (closed symbols) and a G-Rex system (open symbols). Error bars indicate SD (n=4) Comparisons were made by unpaired *t*-test. **(B)** Ratio of nuclei versus cRBC during differentiation in culture dishes or G-Rex (<1 means more cRBC than nuclei). Mean ± SD (unpaired *t*-test, \**P*<0.05, \*\**P*<0.01; n=3-4). **(C)** Enucleation percentages of 10 days differentiated erythroid cultures before and after passage through a leukoreduction filter. **(D)** Cell morphology of the samples analysed in C stained for hemoglobin with benzidine and general cytological stains.

### cRBC resemble mature reticulocytes

Filtered cRBC displayed a spheroid morphology, indicative of a reticulocyte population (Figure 3C). Expression of the major membrane proteins α/ß-spectrin, Band 3, protein 4.1, protein 4.2 and GAPDH were comparable between cRBC from culture dishes and peripheral blood RBC (Figure 4A). Reticulocytes released from the bone marrow have a low deformability, which increases during maturation towards RBC^33^. Both cRBC cultured in culture dishes and the G-Rex system showed a deformability index comparable to late peripheral blood reticulocytes (Figure 4B). Furthermore, cell pellets from cRBC turned dark red and HPLC data showed that cRBC both derived from G-Rex and culture dishes mainly express adult hemoglobin (HbA 1 73.4% G-Rex *vs.* 62% dish; HbA2 2.1% G-Rex *vs.* 1.4% dish) and low levels of HbF (8.8% G-Rex *vs.* 7.1% dish; Figure 4C-D). In addition, hemoglobin oxygen association and dissociation rates of cRBC and peripheral blood RBC were similar at variable oxygen pressure (Figure 4E). Blood group expression of the cRBC was assessed by flow cytometry and compared to the original donor RBC. Blood group expression was in complete agreement with the original donors (Table 1). The functional similarities between cRBC and peripheral blood RBC indicate that the three-stage culture model using culture dishes or a G-Rex bioreactor (Figure 5) yields functional enucleated erythroid cells.

**Figure 4.**
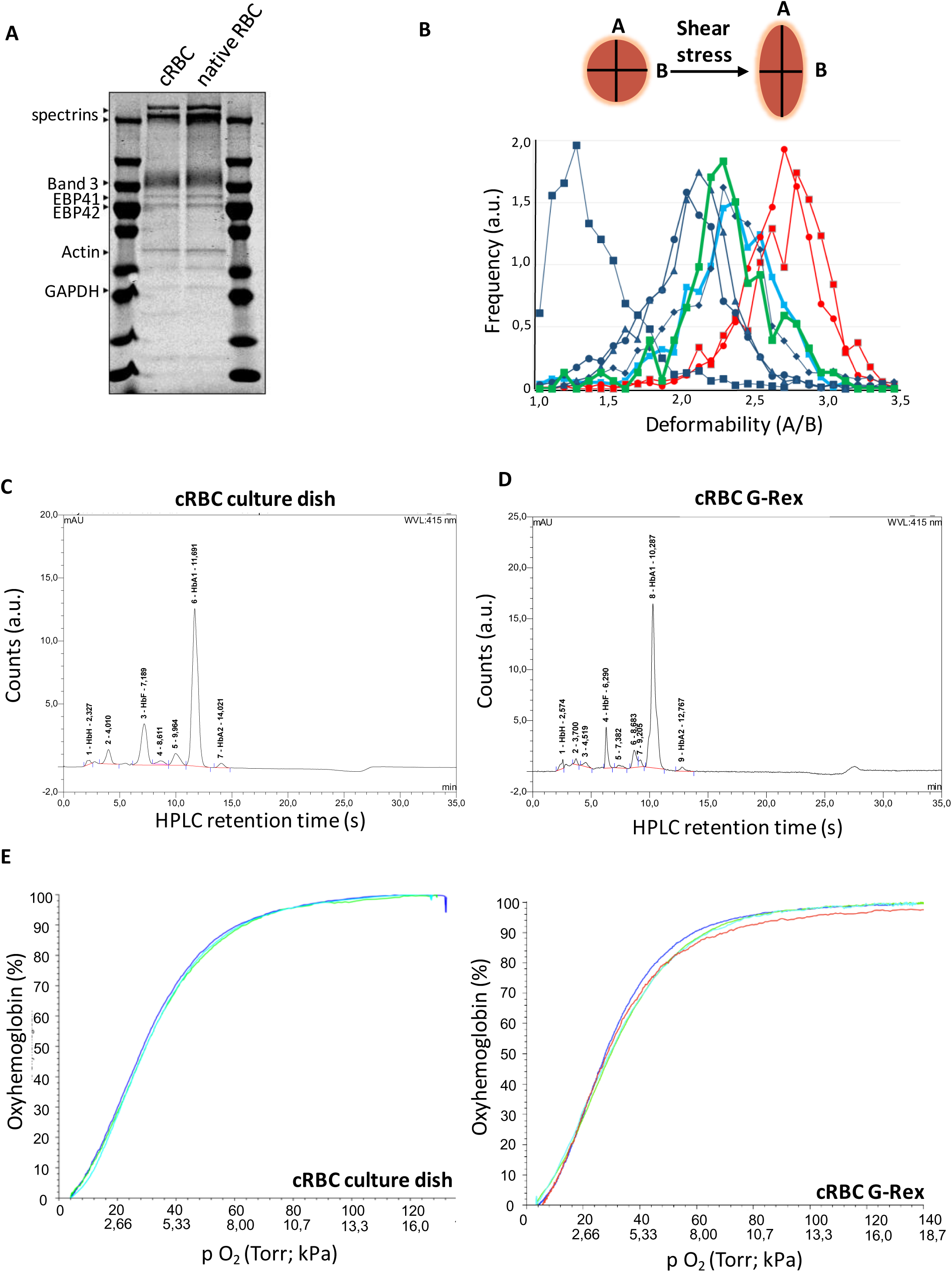
cRBC are highly similar to donor RBC. **(A)** Ghosts from cRBC and peripheral blood RBC were lysed and subjected to SDS-PAGE. Gel was stained with Coomassie and depicts the most abundant proteins in RBC membranes. **(B)** Deformability of cRBC was measured under shear stress by ARCA, which elongates cells and measures length over width as deformability parameter. Progressively maturating reticulocytes were isolated from peripheral blood (in order of maturation, RNA^high^CD71^high^, blue squares; RNA^high^CD71^low^, blue circles; RNA^high^CD71^−^, blue triangles; RNA^low^CD7^−^, blue diamonds; as described^24^) were compared with fully mature RBC (red curves) and filtered cRBC from normal culture dishes (thick green line) or the G-Rex bioreactor (thick blue line). **(C-D)** Expression of hemoglobin variants was determined by HPLC in cRBC in culture dishes before filtering **(C)** or G-Rex after filtering **(D).** Hemoglobin variants are indicated; exact retention time is indicated for each peak. **(E)** Oxygen association and dissociation curve for peripheral blood RBC (teal and red) and cRBC cultured in a G-Rex bioreactor (blue and green). The percentage oxygenated hemoglobin is measured at a gradient of oxygen tension given in Torr (upper line x-axes) and kPa (lower line x-axes).

**Figure 5.**
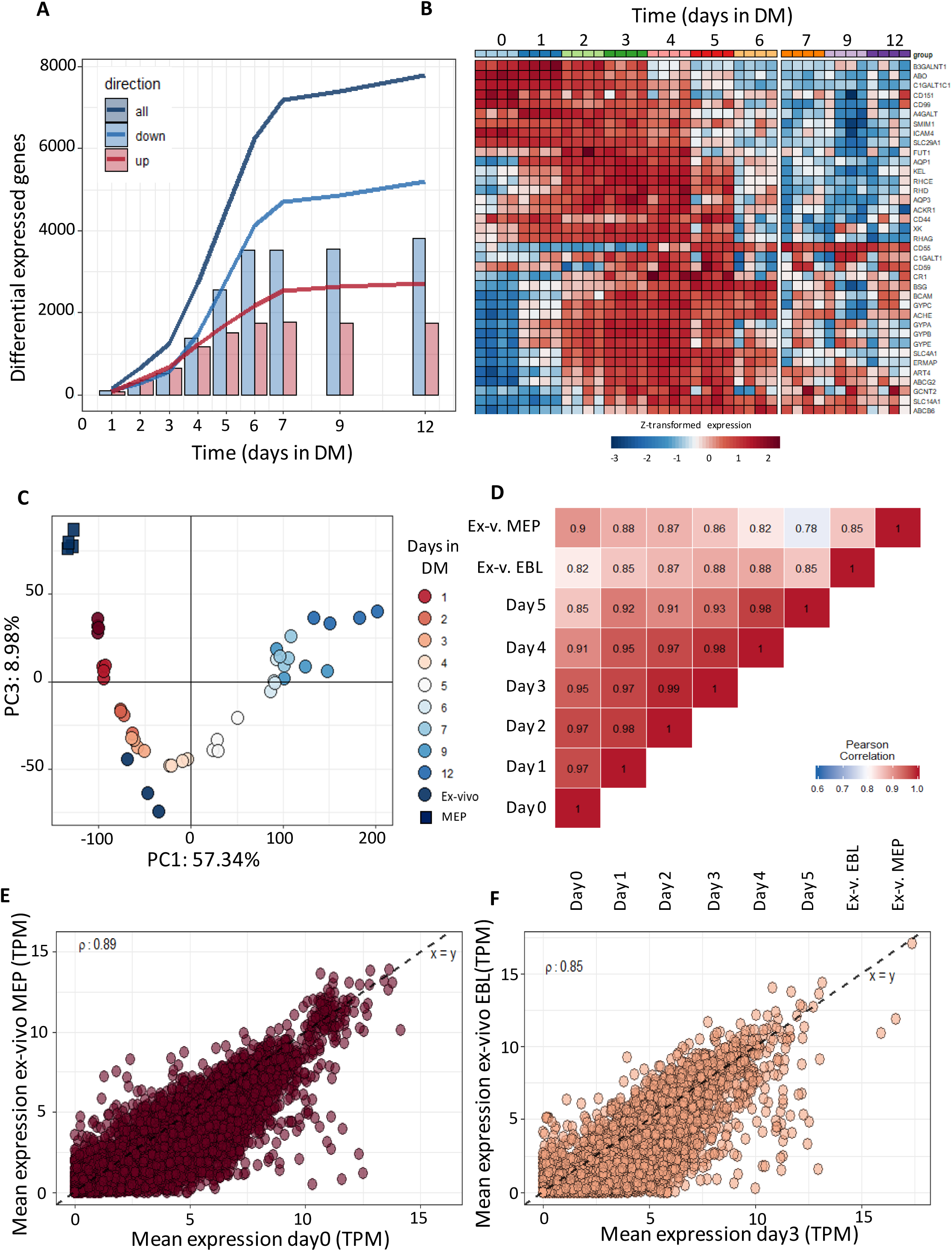
RNA expression profiles during *in-vitro* erythroid differentiation are similar to *ex-vivo* isolated erythroid precursors. RNA was isolated at subsequent days of differentiation to identify changes in the transcriptome (4 independent donors) **(A)** Bars represent the number of transcripts that are up- (red) or downregulated (blue) each day in reference to the start of the culture (FDR < 0.01 and log fold change >2, or < −2). Lines reflect cumulative number of unique genes differentially expressed over the time course. **(B)** Heatmap of z-transformed expression values (log2-CPM) for all genes encoding for blood group antigens and or blood group bearing moieties. **(C)** The transcriptome of MEP (black squares) and CD71^high^CD235^high^ erythroblasts (black circles) isolated from bone marrow was compared to the transcriptome of differentiating cRBC using principle component analysis (PC1 versus PC3). **(D)** A Pearson correlation matrix of all samples used in panel A-C. **(E-F)** Mean transcript levels (transcripts per million mapped reads) were compared between MEP and cRBC day0 (E); and between freshly isolated erythroblasts and cRBC day3 **(F).**

**Table 1:** Blood group analysis of peripheral blood RBC (RBC) and cultured RBC (cRBC) using flow cytometry.

| Blood group | RBC<br>1 | RBC<br>2 | RBC<br>3 | cRBC<br>1 | cRBC<br>2 | cRBC<br>3 |
| --- | --- | --- | --- | --- | --- | --- |
| **A** | - | - | + | - | - | + |
| **B** | - | - | - | - | - | - |
| **RhD** | + | + | - | + | + | - |
| **RhE** | - | + | - | - | + | - |
| **RhC** | + | - | - | + | - | - |
| **Rhe** | + | + | + | + | + | + |
| **Rhc** | + | + | + | + | + | + |
| **K** | - | - | - | - | - | - |
| **k** | + | + | + | + | + | + |
| **Kp-a** | - | - | - | - | - | - |
| **Kp-b** | + | + | + | + | + | + |
| **Lu-a** | - | - | - | - | - | - |
| **Lu-b** | + | + | + | + | + | + |
| **P1** | + | - | + | + | - | + |
| **S** | + | - | - | + | - | - |
| **s** | + | + | + | + | + | + |
| **Fya** | + | - | + | + | - | + |
| **Fyb** | - | + | + | - | + | + |
| **Jka** | - | + | - | - | + | - |
| **Jkb** | + | - | + | + | - | + |
| **M** | + | - | + | + | - | + |
| **N** | + | + | + | + | + | + |

### Terminal differentiation of erythroblasts to enucleated reticulocytes completely changes the transcriptome

Differentiation of non-hemoglobinized erythroid progenitor cells to functional enucleated cells involves significant changes in morphology, cell volume and protein content. Identification of regulatory processes is crucial to enhance and optimize the production and the yield of *in-vitro* erythropoiesis. A comparison of the *in-vitro* transcriptome to existing databases of *in-vivo* transcriptomes allows to benchmark the *in-vitro* differentiation process. Differentiation was started from CD117^+^CD71^+^CD235^−/dim^CD44^high^ early erythroblast populations (phase III, day 0) from four distinct donors and daily RNA samples were collected until terminal enucleation state at day 12 (95%; Changes in surface marker expression in Figure S2). In reference to the early erythroblast population (day 0) 75% of the analyzed transcripts changed during differentiation (7792 transcripts with a FDR < 0.01 and fold-change > 4) with 5189 down regulated, 2726 upregulated and 124 that were transiently up- or downregulated (Figure 5A and Table S2). These major transcriptome alterations occurred primarily during the first 6 days (phase III). The expression of transcripts encoding proteins crucial for the function and immunological properties of RBC can be scrutinized. Figure 5B shows that RNA expression of different blood group antigen bearing proteins from these donors is differentially regulated over time. In addition, globin subunit expression dynamics indicated rapid hemoglobinization during the first 72 hours of differentiation coinciding with increased CD71 expression. Note that expression of beta globin subunits is significantly higher compared to gamma globins. *cKIT* RNA was rapidly downregulated and *CD235* (GPA) rapidly upregulated in agreement with flow cytometry results (Figure S4A). In line with the major transcriptional changes over the course of differentiation, principal component analysis (PCA) captured ~57% of variance in the 2000 most variable genes. The first component associated with differentiation progression, sequentially grouping samples from subsequent days (Figure 5C). Cultured RBC showed good functional and morphological correspondence with *in-vivo* RBC (Figure 4). This raised the question how the cultured cells would compare to *ex-vivo* cells at the transcriptome level. Comparing published transcriptomes of bone marrow megakaryoid/erythroid progenitors (CD38^+^CD34^+^CD10^−^CD45RA^−^CD123^−^CD90^−^) and CD71^+^CD235^+^erythroblasts^34^ to *in-vitro* cultured erythroid cells revealed that the *ex-vivo* MEP grouped before the sequence of cultured cells, while the CD71^+^CD235^+^ grouped along with the cultured cells, with the same marker expression (Figure S4B). Note that the *in-vitro* cultures showed little variation between donors indicating a reproducible synchronized differentiation process. *Ex-vivo* cells showed relatively more variation in the PCA, which may be a consequence of the more broadly expressed erythroid surface markers used to isolate these cells. The major differences between cultured and *ex-vivo* cells accounted for 18% of the variance (Figure S5A). Still, Pearson correlation between samples using all differentially expressed genes indicated that *ex-vivo* MEP were most similar to erythroblasts cultured under expansion conditions (d0) whereas *ex-vivo* CD235^+^CD71^+^ erythroblasts were most similar to d3 differentiated erythroblasts (ρ = 0.89 and ρ = 0.85 respectively, Figure 5D). Direct comparison of RNA expression between the *ex-vivo* MEP and do erythroid cells, and of the CD235^+^CD71^+^ *ex-vivo* erythroblast and d3 differentiated cells showed comparable transcript levels (Figure 5E-F). Overall, the main difference between cultured and freshly isolated cells originates from genes that were predominantly expressed at increased levels in cultured cells compared to *ex-vivo* cells (Table S3). Of note, transcripts with increased expression in cultured cells were further downregulated upon differentiation progression in line with decreased expression observed in the more asynchronous *ex-vivo* erythroid cells (Figure S4B). Removing the low expression filter also revealed a set of 380 genes expressed at lower levels in cultured cells, that consisted of pseudogenes and transcripts encoding mitochondrial or ribosomal proteins (Table S4), probably reflecting a difference in technical processing of samples. The similarities in the transcriptome of cultured cells and the related stage *in-vivo* indicates that close recapitulation of transcriptional changes is at the basis of the functional and morphological characteristics of the cultured erythroid cells.

## Discussion

Widescale clinical application of cRBC is faced by several constraints, like the inability to generate large cell numbers that are required, the high costs of ill-defined media and the low yield of enucleated, biconcave cRBC^35^. The culture protocol presented here challenges these constraints by boosting advances with respect to high enucleation rates, matching erythroid characteristic at different levels, redefining medium composition and maximum expansion without the necessity to first isolate CD34^+^ cells (Figure 6). Using a defined medium and exploiting stress erythropoiesis we achieve 10^7^-fold erythroblasts expansion within 26 days. Despite starting from total PBMC, pure erythroid cultures expressing CD71 and CD235a^dim^ are obtained validating the GMP-grade medium yielding similar cell numbers as previously reported using commercial Stemspan media^17,18^. Using total PBMC not only allows outgrowth of all CD34^+^ progenitors, but also CD34’ hematopoietic progenitors present in blood that have the capacity to differentiate towards erythroid cells^17,18^. Glucocorticoids are essential for stress erythropoiesis in the mouse and synergy of glucocorticoids with EPO and SCF induces erythroblasts proliferation whilst inhibiting differentiation^7,23,32,36,37^. The addition of glucocorticoids also supports erythropoiesis by differentiating peripheral blood monocytes or CD34^+^ cells to erythroid supporting macrophages during culture from PBMC, further increasing the erythroid yield^19,38^. In contrast to our plasma/serum-free expansion, many large-scale red cell culture protocols use glucocorticoid agonists in combination with serum and/or plasma during expansion. Here, we showed that the addition of plasma causes premature differentiation of erythroblasts also in the presence of glucocorticoids^9,10,13,39,40^. Increased spontaneous differentiation upon addition of plasma during erythroblast expansion may be due to additional growth factors or other plasma components. Optimal expansion in absence of plasma/serum is important to reach the amount of cRBC required for transfusion, but also to establish erythroid cultures from small blood aliquots of specific anemic patients. We have recently shown that 3ml of blood is sufficient to culture enough cells to reprogram erythroblasts to induced pluripotent stem cells (IPSC)^41^. In addition to direct use, the expanded erythroblast can be frozen and defrosted without loss of expansion potential (data not shown), similar to what we previously reported for starting cultures from frozen PBMCs^42^. This introduces flexibility concerning actual production of products. For instance, we have previously shown that cryo-preserved patient-derived erythroblasts can be reprogrammed to İPSC^43–45^.

**Figure 6.**
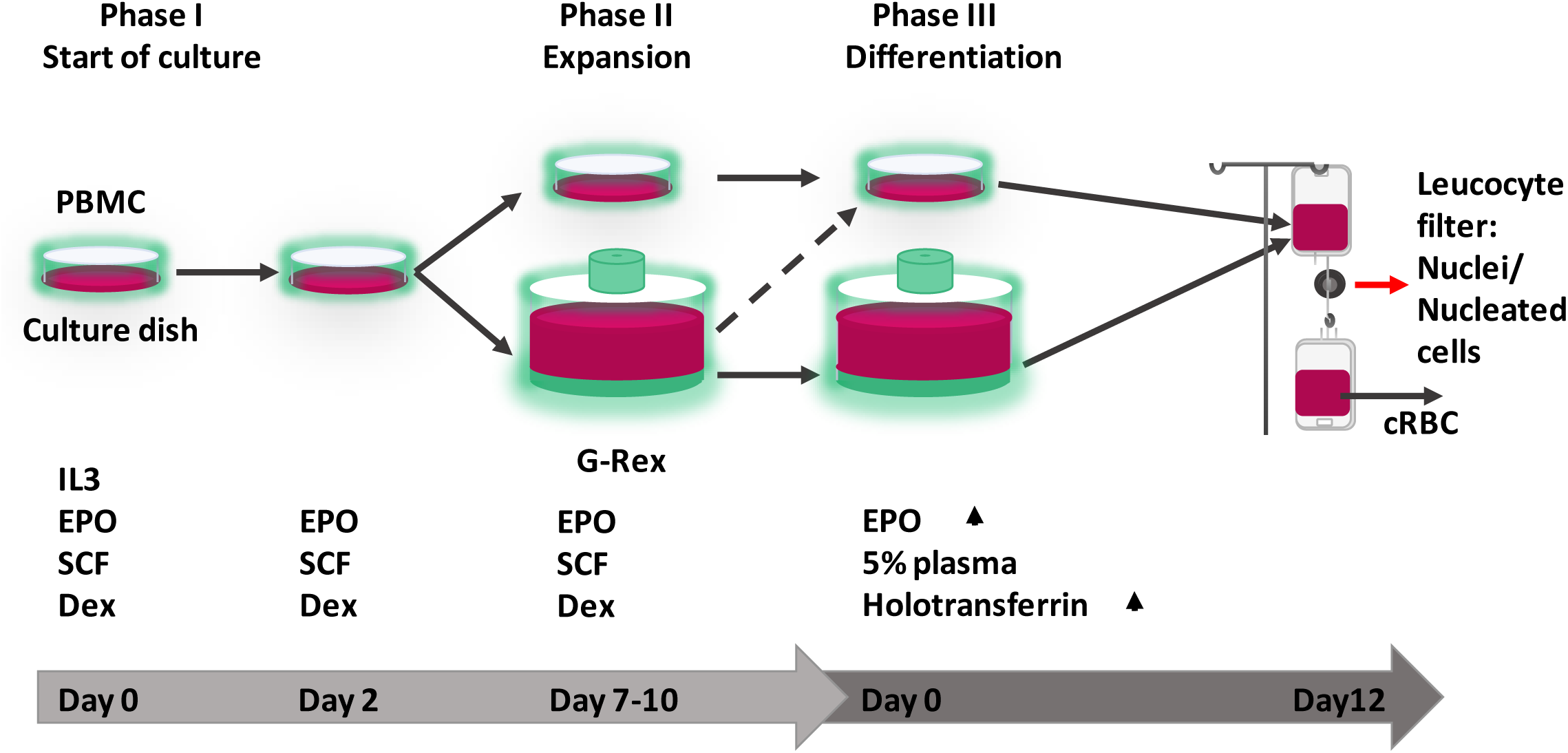
Progression of erythroid cells during culture. Overview of the three-phase erythroid culture system using culture dishes or a G-Rex bioreactor. A pure erythroblast culture is established in dishes from PBMC during the first 7 days. From day 7 erythroblasts are expanded in G-Rex or in dishes. When transferred from expansion to differentiation medium, erythroblasts in dishes or in G-Rex can mature to enucleated cells. The small remnant of nucleated cells and nuclei (pyrenocytes) can be removed by passage through a leucocyte filter. The growth factors and hormones used in culture are indicated at the lower half of the graph.

The use of adult PBMC also facilitates the availability of starting material and introduces the possibility to culture autologous cRBC. This is important considering alloimmunization caused by blood group mismatches and matching from cord blood derived RBC may be complicated. In addition, with the advances of using therapeutic amounts of RBC loaded with specific cargo, using the patient’s own cells is also less complicated compared to unmatched blood, cord blood or immortalized cell lines like human IPSC or immortalized erythroblasts.

The culture process was compatible with up-scaling for clinical studies and applications using a G-Rex bioreactor. The G-Rex bioreactor allowed for 3×10^6^ fold expansion per PBMC. Remaining nuclei and enucleated cells after differentiation could be efficiently removed using a leukoreduction filter, as generally employed by bloodbanks, to obtain a pure cRBC population. The defined IMDM-based culture medium, termed Cellquin, solely contains GMP grade components and finds its basis in HEMA-def^7^. Knowing the exact concentrations of all components within Cellquin now allows to quantitatively track erythroid requirements by combining the transcriptome/proteome with metabolomics. This may help to culture cells at higher densities and to cater specific media components exclusively erythroid need, leading to considerable cost-reduction and aiding upscaling, currently, the costs of Cellquin is lower compared to commercially available media, while performing at least similar.

One Cellquin component paramount to its effectiveness is albumin. We observed that the isolation and manufacturing process of human albumin critically influences the erythroid expansion potential. Using ultraclean, detoxified, or recombinant additive-free HSA significantly increased the erythroid expansion potential. Albumin binds substances including proteins, metabolites and fatty acids, including toxins, drugs and other therapeutics^46–48^. Replacing ultra-pure cHSA in EM by Albuman^®^ reduced erythroblast expansion potential, which could be reverted by charcoal and ion exchanger treatment of Albuman^®^. Interestingly, the process to manufacture Albuman^®^ includes a saturation step to restrain the albumin binding potential, rendering it mostly inert. This suggests that the binding and/or transport function of albumin is important to ensure continued erythroblast expansion.

The technical improvements to the culture protocol result in excellent yield of *in-vitro* cultured erythroblasts combined with >90% enucleation, adult hemoglobin expression, correct blood group expression, deformability and oxygen saturation dynamics similar to donor peripheral blood enucleated cells. In addition, we present the first transcriptomic analysis from erythropoiesis originating from stress-erythropoiesis cultures. Comparison of this dataset to datasets from *ex-vivo* erythroid cells shows that, next to similarities in RBC characteristics, the cRBC are comparable to similar *ex-vivo* cells at transcript level. These observations together, make the transcriptome dataset provided here a valuable resource to address erythroid regulatory mechanisms.

In 2011, Timmins et al. demonstrated an ultra-high yield of cRBC with >90% enucleation from cord blood-derived CD34+ cells in the absence of plasma^15^. This reported erythroid yield was similar to our serum/plasma-free adult PBMC-derived erythroid expansion. However, >90% enucleation during terminal differentiation using our differentiation protocol (stage III) was only recapitulated in presence of plasma. Where, adding 5% Omniplasma increases enucleation from 20-25% to more than 90% (data not shown).Whether this reflects a difference in cord blood versus adult erythroid cultures remains unknown but important to investigate along with identifying the components in plasma that are key to this increased enucleation in our system.

Combined the advances of the presented protocol facilitate both easy implementation in other laboratories and study of erythropoiesis in healthy individuals or in patients of which limited sample volumes are available. It adds to the feasibility of using adult peripheral blood as starting material for cRBC cultures, which is an important step towards precision medicine. For example, in using custom engineered cRBC as cargo vesicles for drug delivery. In addition, the synchronized cultures contain young reticulocytes that have a theoretical lifespan of about 120 days. Chronic anemia patients that receive blood transfusions every two months may benefit from transfusions with *in-vitro* cultured long-lived RBC, potentially increasing the time between transfusions and thereby reducing the costs. Currently we are working towards a clinical trial that will allow to test the *in-vivo* lifespan of transfused PBMC-derived cRBC.

## Author Contributions

PB, MT-V, ES, EH, SH, EO, EV and AV performed the experiments. EA, EH, and PB designed the experiments, analyzed the data and wrote the manuscript. JM performed the RNA isolation and sequencing. ML contributed to the experiment design and writing of the manuscript. The manuscript was critically revised by all authors. The authors declare no competing financial interests.

## Acknowledgments

We are grateful to Wilson Wolf Manufacturing (Saint Paul, MN, USA) for providing the G-Rex bioreactors. We would like to thank Rob van Zwieten, Martijn Veldthuis and Jeffrey Berghuis (Sanquin, dept. Blood Cell Research) for the technical assistance and data acquisition regarding the HPLC data and deformability assays, and the Central Facility of Sanquin for their assistance regarding flow cytometry analysis. This work was supported by grants from The Netherlands Organization for Health Research and Development (ZonMw-TOP, 40-00812-98-12128; SH and ZONMW-TAS, 40-41400-98-1327; PB, MT-V, ES), from the Landsteiner Foundation for Blood Transfusion Research (LSBR:1141; EA, EH),; and by Sanquin (PPOR:15-30; AV).

## Supplementary material

Additional figures can be found in supplementary information. RNA-sequencing data has been submitted to NCBI’s Gene Expression Omnibus and is available under accession number: GSE124363.

