## Supplemental material for "Large-scale *in-vitro* production of red blood cells from human peripheral blood mononuclear cells"

**Supplementary methods and material**

**Cell culture using different sources of human albumin**

EM medium was supplemented with human bovine albumin (HSA) from different sources: Albuman® (Sanquin, Amsterdam, The Netherlands), detoxified HSA^1^, recombinant HSA (Akron Biotech, Boca Raton, FL, USA), or AB+ pool plasma (Sanquin) at indicated concentrations as specified in the figure legends. In addition, the commercial medium Stemspan (Stem Cell Technologies, Germany) was used ^2^.

**RNA-sequening**

Erythroblasts were allowed to differentiate to enucleated reticulocytes as described in culture protocol and summarized in figure 6. Cells were collected daily; RNA was isolated using Trizol (Invitrogen Technologies, Carlsbad, CA) and amplified and rRNA depleted using HyperPrep Kit with RiboErase (KAPA Biosystems, Pleasanton, CA, USA) as described by manufactures.

Samples were sequenced at 30 million 75bp paired-end reads. After quality control with FastQC, read were mapped to genome ChGR38.v85 using STAR ^3^. Lowly expressed genes were filtered from count data, after correction for library size (counts per million mapped reads < 3). Subsequent differential expression analysis was performed with EdgeR using a paired design for individual donors and quasi-likelihood F-test to test expression at each day compared to day 0 ^4^. The differential expression threshold was set over 4-fold difference and an FDR < 0.01.

The comparison of cRBCs to ex-vivo cells was performed on samples aligned to ChGR38.v85 with Salmon ^5^. This aligner allows for direct mapping to transcript isoforms, including these in the comparison while facilitating normalization to transcript length prior to plotting. Counts were normalized for transcript length and library size, as transcript per million mapped reads (TPM) and were plotted using ggplot2 package in R ^6^.

**Culture filtration to obtain reticulocytes**

CRBC were purified using a 60ml neonatal leucoreduction filter (Haemonetic BV, Breda, The Netherlands). The filter was primed with IMDM containing 4% Omniplasma. Cultured cells were filtered followed by a 2 times wash with 25ml priming medium. CRBC were resuspended in IMDM containing 4% Omniplasma. Cells were centrifuged at 1600rpm for 5 min in order to obtain packed red blood cells.

**Flow cytometry**

Antibodies used: Acris (Herford, Germany): anti-CD235a (FITC 1:400; PE 1:400); BD Biosciences: anti-CD117 (PE 1:50), anti-CD235a (PE 1:150); eBioscience (Vienna, Austria): anti-CD44 (APC 1:150); Miltenyi Biotec: anti-CD71 (APC 1:100; VioBlue 1:200), anti-CD235a (VioBlue 1:200); and Life technologies (Carlsbad, CA, USA): anti-CD117 (FITC 1:100). The following antibodies were used for blood group analysis: IBGRL research products (Bristol, UK): anti-A (Bric145; 1:1000), anti-B (BGRL2; 1:500), anti-M (clone 6A7; 1:5000), anti-N (clone BRIC157; 1:1000); Sanquin Products (Amsterdam, The Netherlands): anti-RhC (clone MS24; RBC 1:1000, cRBC undiluted), anti-Rhc (clone MS35; 1:200), anti-RhE (clone MS260; RBC 1:80, cRBC undiluted), anti-Rhe (clone MS21/63; RBC 1:100, cRBC undiluted), P1 (clone P3NIL100; 1:500), Kp-a (undiluted), kp-b (undiluted), Jk-a (MS15; undiluted), Jk-b (MS8; undiluted), Fy-a (P3TIM; undiluted), Fy-b (undiluted), Lu-a (undiluted), Lu-b (undiluted); Merck-Millipore (Darmstadt, Germany): anti-K (MS59; 1:50), anti-k (P3A118OL67; 1:100), anti-s (P3Y326Bn5; 1:400). Ortho Clinical Diagnostics (Buckinghamshire, UK): anti-S (MNS3; undiluted). Anti-A, B, N, M were detected with an anti-mouse IgG FITC-labeled antibody (Life Technologies; 1:200) was used and anti-C, c, E, e, Jk-a, Jk-b, and P1 with an anti-human IgM PE-labeled antibody (Southern Biotech, Birmingham, AL; 1:500). The remaining blood groups were detected with an anti-human IgG PE-labeled antibody (Southern Biotech; 1:200).

Supplementary Figure Legends

**Figure S1. The presence of plasma/serum affects erythroblast expansion. (A)** Representative dot plots of erythroblasts at day 12 of culture in EM in the presence of ultra-clean HSA (cHSA) or EM in which cHSA was replaced by plasma, Albuman® or detoxified HSA (dHSA) corresponding to Figure 1A (n=3-4). Note that the percentage of immature erythroblasts (P1) is 3-4 times higher in cHSA and dHSA compared to plasma and Albuman®. This suggests that cHSA and dHSA prolong the immature state of erythroid cultures. **(B)** Erythroblasts cultured in EM for 18 days in the presence of cHSA, dHSA or recombinant HSA (rHSA) showed a similar expansion potential. Mean ± SD (two-way ANOVA; n=2). **(C)** Erythroblasts at day 7 and 19 of expansion in Stemspan, as described previously^2^.

**Figure S2. Marker expression during erythroblast differentiation in culture dishes. (A)** Erythroblasts were cultured in EM for 8 days and subsequently differentiated for 12 days in DM. Representative overlaying histograms showing the FSC-A during differentiation and the expression levels of CD71, CD235a, CD117 and CD44 analyzed by flow cytometry. **(B)** Graphs corresponding to panel A showing the mean fluorescence intensity (MFI; n=4). **(C)** Lines depicts RNA expression over days of differentiation (log2 transformed CPM) for differentiation markers CD71 (transferrin receptor, TFRC) and CD235 (glycophorin A, GYPA) along with c-kit(SCF receptor, KIT) adult globin subunits (HBB and HBA2) and fetal globin subunit (HBG2).

**Figure S3. Enucleation of cRBC derived from culture dishes or a G-Rex bioreactor**. **(A)** Graph showing the enucleation of erythroid cells during differentiation after supplementation into DM to initiate differentiation in a G-Rex bioreactor, compared to culture dishes corresponding to Figure 2E. Mean ± SD (paired *t*-test, **P<*0.05, ***P<*0.01, ****P<*0.001; n=4). **(B)** Erythroblasts expanded and differentiated in culture dishes with 90% (unwashed) or 100% (washed) replacement of expansion medium before differentiation medium. Mean ± SEM (paired *t*-test, ***P<*0.01; n=4). **(C)** Two days prior to differentiation in DM, SCF was either removed from or maintained in EM and enucleation was analyzed after 12 days of differentiation. Mean ± SEM (paired *t*-test, *****P<*0.0001; n=4). **(D)** Two days prior to differentiation 1000nM or 10nM dexamethasone was added to the expansion medium. Expansion medium was replaced with differentiation medium and cells were analyzed for enucleation after 12 days. Mean ± SEM (paired *t*-test; n=4). **(E)** Representative dot plots of erythroblasts during differentiation in DM in culture dishes or G-Rex based on the expression of DRAQ5.

**Figure S4. Variation between cRBC and ex-vivo cells. (A)** Plot of cRBC and ex-vivo cells on first and second component of PCA. **(B)** Line plot shows expression levels for transcripts higher expressed in D3 cRBC compared to ex-vivo cells. **(C)** Correlation matrix and expression plots without filtering lowly expressed genes.

6.	H W. ggplot2: Elegant Graphics for Data Analysis. New York: Springer-Verlag; 2016.
