## Supplementary figures and images for "Large-scale *in-vitro* production of red blood cells from human peripheral blood mononuclear cells"

### Supplemental figures

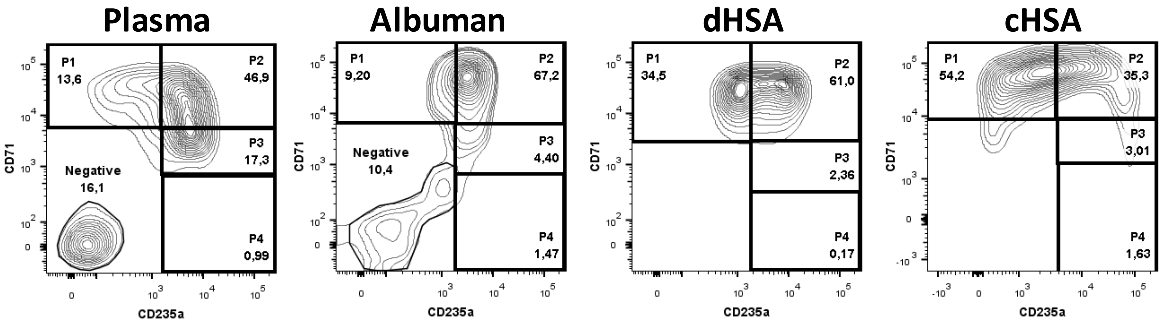

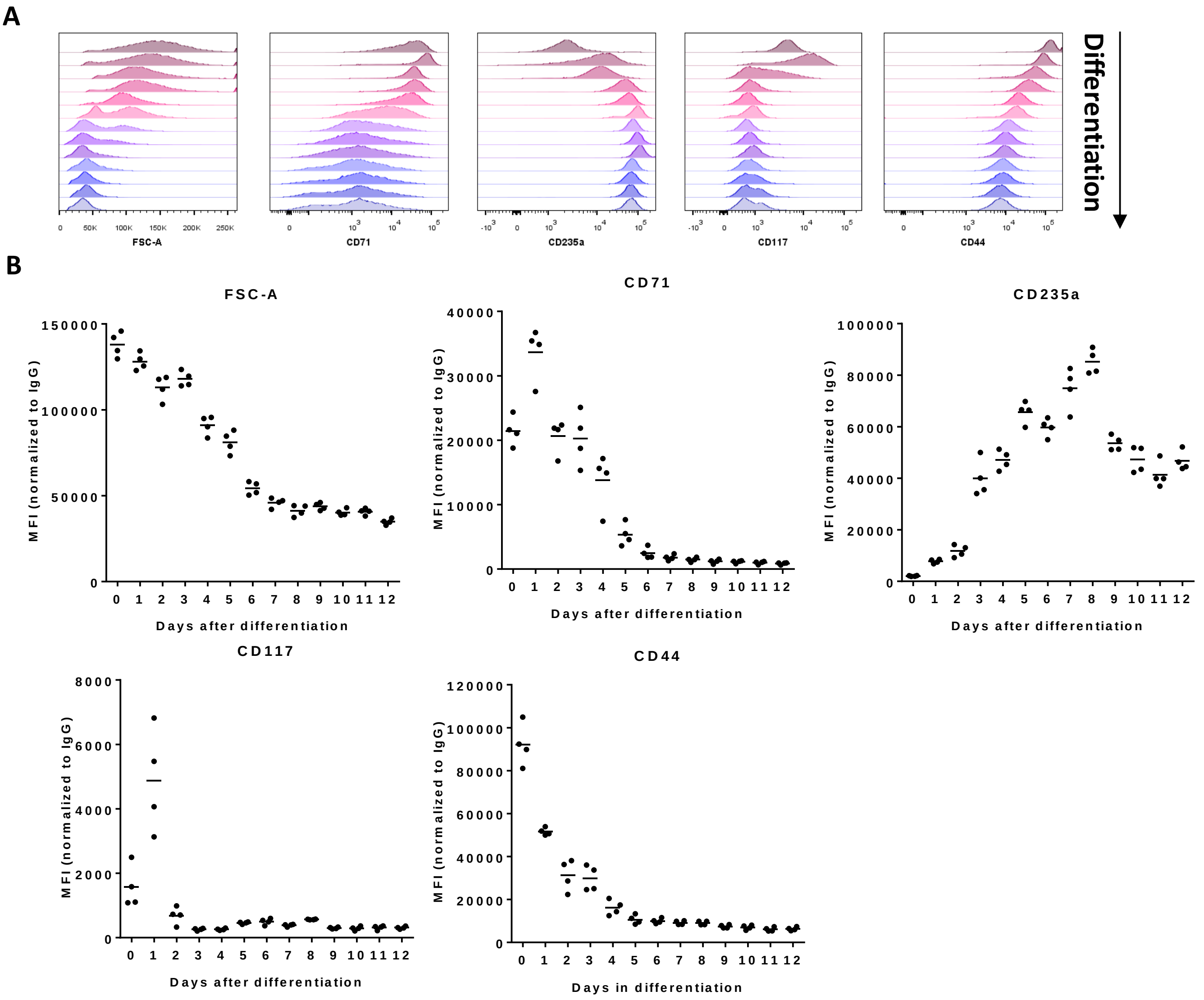

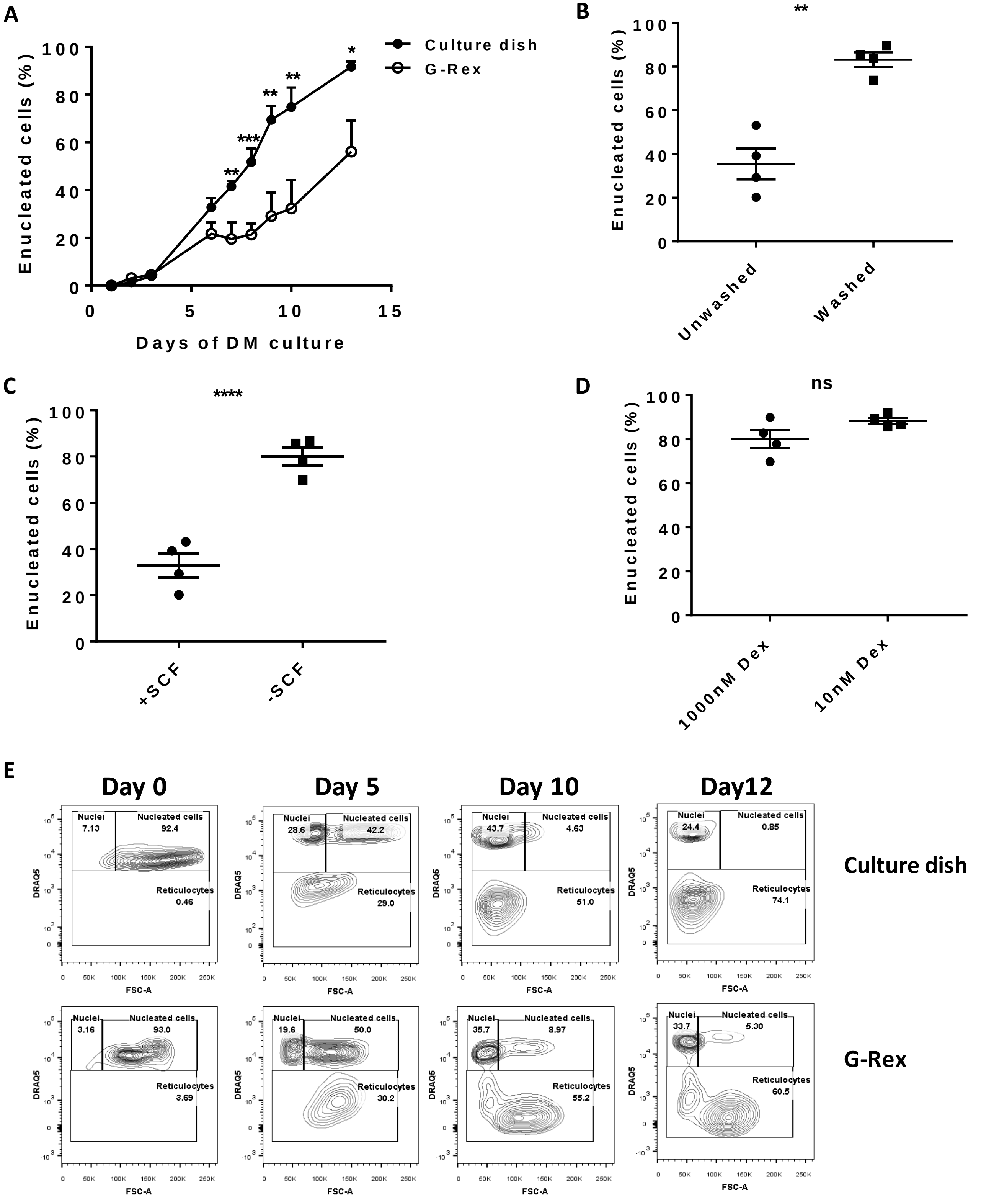

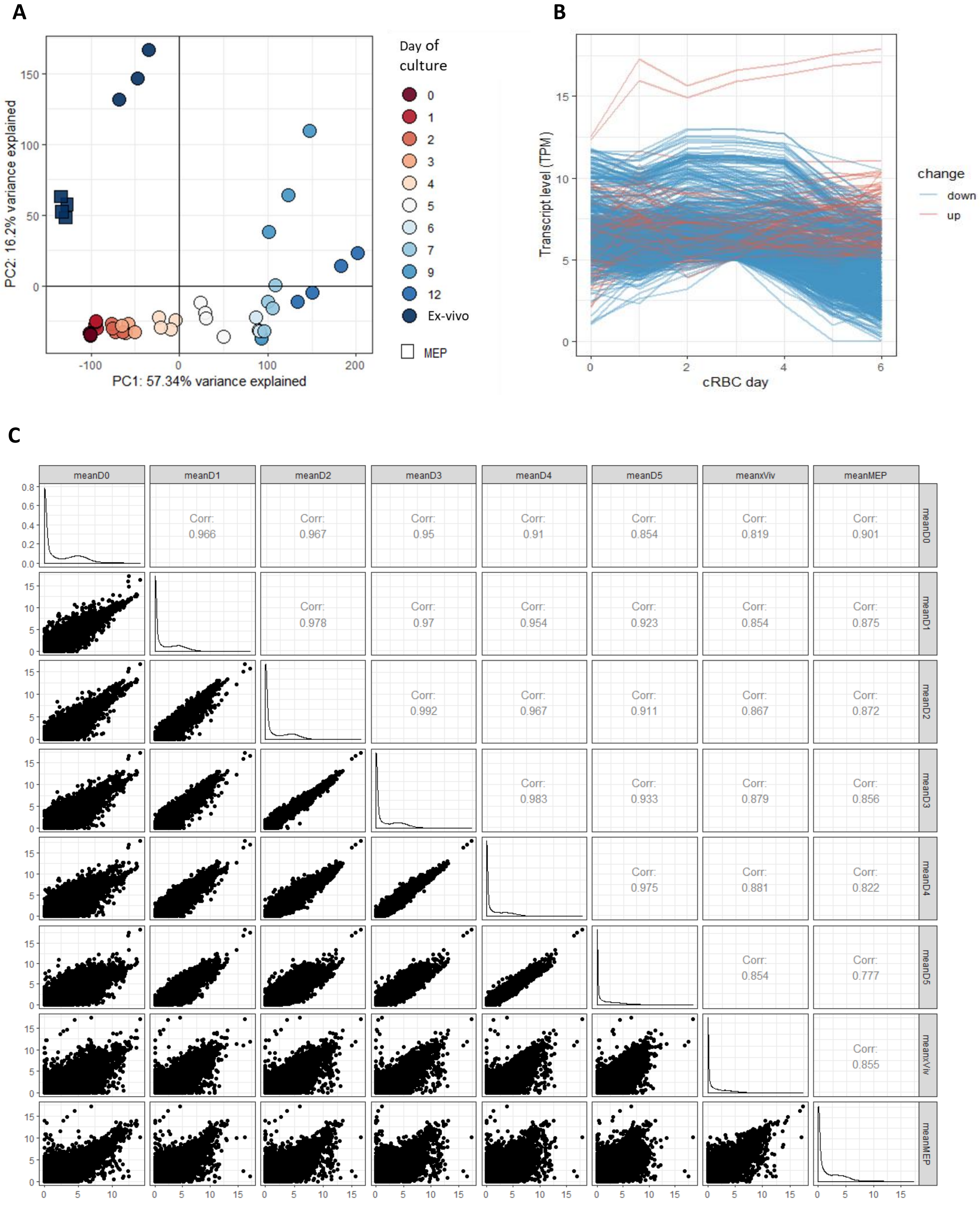

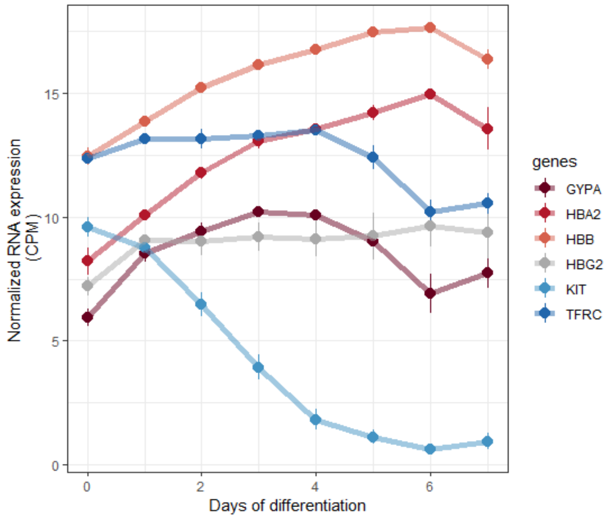

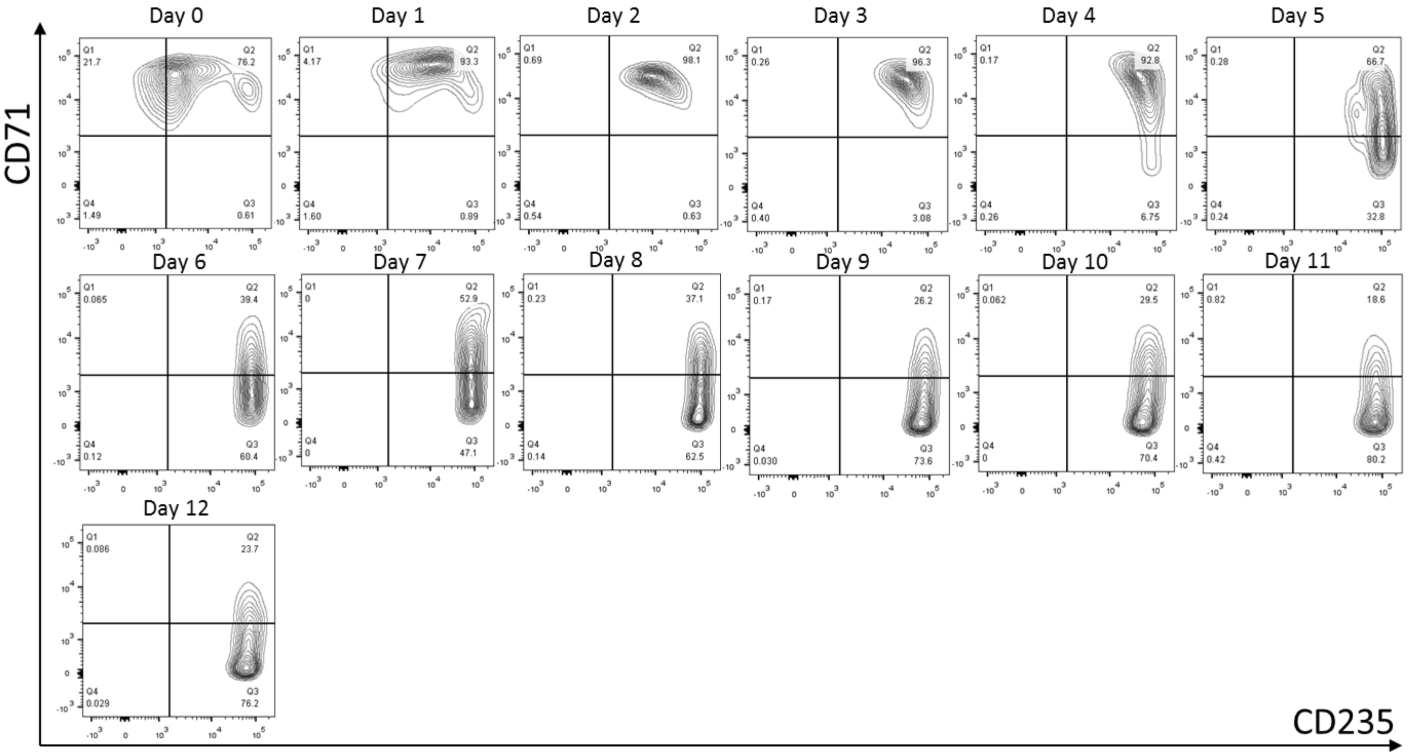
